# Transmembrane protein 184B (TMEM184B) promotes expression of synaptic gene networks in the mouse hippocampus

**DOI:** 10.1101/2022.05.27.493793

**Authors:** Elizabeth B. Wright, Erik G. Larsen, Cecilia M. Coloma-Roessle, Hannah R. Hart, Martha R.C. Bhattacharya

## Abstract

In Alzheimer’s Disease (AD) and other dementias, hippocampal synaptic dysfunction and loss contribute to the progression of memory impairment. Recent analysis of human AD transcriptomes has provided a list of gene candidates that may serve as drivers of disease. One such candidate is the membrane protein TMEM184B. To evaluate whether TMEM184B contributes to neurological impairment, we asked whether loss of TMEM184B in mice causes gene expression or behavior alterations, focusing on the hippocampus. Because one major risk factor for AD is age, we compared young adult (5-month-old) and aged (15-month-old) wild type and *Tmem184b*-mutant mice to assess the dual contributions of age and genotype. TMEM184B loss altered expression of pre- and post-synaptic transcripts by 5 months and continued through 15 months, specifically affecting genes involved in synapse assembly and neural development. Wnt-activated enhancer elements were enriched among differentially expressed genes, suggesting an intersection with this pathway. Few differences existed between young adult and aged mutants, suggesting that transcriptional effects of TMEM184B loss are relatively constant. To understand how TMEM184B disruption may impact behaviors, we evaluated memory using the novel object recognition test and anxiety using the elevated plus maze. Young adult *Tmem184b-*mutant mice show normal object discrimination, suggesting a lack of memory impairment at this age. However, mutant mice showed decreased anxiety, a phenotype seen in some neurodevelopmental disorders. Taken together, our data suggest that TMEM184B is required for proper synaptic gene expression and anxiety-related behavior and is more likely to be linked to neurodevelopmental disorders than to dementia.

## Introduction

Alzheimer’s Disease (AD) is a devastating neurodegenerative disease for which we have a paucity of treatment options. In 2022, it was estimated that 10.7% of adults over 65 have Alzheimer’s-induced dementia [1]. The burden on patients, their families and health systems is expected to grow significantly in the next 20 years; this trend is already apparent [2]. Alzheimer’s Disease causes the accumulation of amyloid beta plaques and neurofibrillary tangles in diverse regions of the brain [3]. Many clinical trials have focused on the depletion of amyloid as a key therapeutic strategy. However, trials in which amyloid plaques are successfully diminished have not produced a reduction or delay in cognitive impairment in trial participants [4]. This highlights the need for the identification of new targets that may contribute to early cognitive impairment and that may offer hope for alternative strategies to slow the progression of Alzheimer’s Disease.

Large-scale human genome-wide association and RNAseq analysis to identify network drivers of disease has yielded many plausible candidates that could contribute to AD [5–7]. With these candidates in hand, the task for the AD field now shifts to identifying which candidates cause bona fide alterations in relevant gene expression pathways or in behaviors associated with AD. Through recent RNAseq analyses, the transmembrane protein TMEM184B was predicted as a candidate protein that could promote AD progression [8]. TMEM184B is a 7-pass transmembrane protein expressed broadly in the nervous system and is thought to play a role in neuronal excitability, synaptic structure, and expression of key developmental and adult pathways involved in neuronal differentiation [9, 10]. Using analysis of the ROSMAP and MAYO clinic human cohorts, TMEM184B was predicted to drive AD-associated gene expression patterns in the dorsolateral prefrontal cortex and temporal cortex, respectively [8]. However, knockdown of TMEM184B in human induced pluripotent stem cell-derived neurons did not directly affect Aβ levels [5], suggesting that this gene may play a more indirect role.

In this study we sought to directly test the hypothesis that TMEM184B regulates gene networks that could contribute to AD, using a mutant mouse model in which the *Tmem184b* gene is disrupted [11]. We compared how TMEM184B-regulated gene networks are influenced by aging, focusing on the hippocampus as a key location involved in memory. We identified impairments in pathways controlling synapse assembly and function in *TMEM184B*-deficient hippocampus. We also evaluated memory and anxiety behaviors in these mice. Our results identify a clear effect of TMEM184B on the expression of synaptic gene networks at both young adult and older ages, specifically on proteins known to promote proper synaptic development and function. While young adult *Tmem184b-*mutant mice do not have impaired memory, we uncovered an unexpected effect of TMEM184B loss on anxiety behavior. Our work places TMEM184B into a gene regulatory network that could impact synaptic connectivity and, rather than cause dementias, may instead be linked to neurodevelopmental disease.

## Materials and Methods

### Mouse Strains

Mice used in this study have a gene-trap insertion originally created by the Texas A&M Institute for Genomic Medicine allele Tmem184bGt^(IST10294F4)^ on the C57BL/6 background. Mice containing this insertion have been previously characterized to have very low (<5% of wild type) *TMEM184B* mRNA expression by both qPCR and RNAseq [10, 11]. C57BL/6 mice bred in house (littermates when possible) were used as controls.

### Hippocampal RNA Isolation

Mice were humanely euthanized with carbon dioxide, and hippocampi were removed within 10 minutes of euthanasia and immediately frozen for subsequent RNA isolation. Total RNA was extracted using Trizol (Invitrogen) using the manufacturer’s protocol. Following initial quality check via Nanodrop analysis, total RNA samples were frozen and shipped to Novogene (Sacramento, CA) for sequencing. Quality control checks were done by Novogene to ensure that sequenced samples had high RNA quality. Sequencing was performed using paired end 150bp reads, 30 million reads/sample, on the Illumina platform. Raw reads data were provided by Novogene. RNAseq data includes the following mice by age and sex: 4 5-6 month wild type (1F/3M), 3 5-6 month mutant (1F/2M), 3 15 month WT (3F, excluded 1 M sample due to poor quality), 3 15-month mutant (1F/2M).

### Data Processing and Statistical Analysis

Alignment to the mouse genome (mm10.0) was done using Salmon and run on the High Performance Computing cluster at the University of Arizona. Differential gene expression analysis was done using DESeq2 on Galaxy servers [12]. PCA plots were generated by Galaxy’s DESeq2 default settings [13], or done in R using the stats R package.

For Gene Ontology (GO) analysis, Panther pathway analysis, and KEGG pathway analysis[14–16], ribosomal genes were manually removed from the lists prior to submission. GO analysis [17] was performed with a background list of genes consisting of the full list of mapped genes identified by pseudoalignment via the Salmon algorithm from the datasets used for each comparison. KEGG analysis was performed in R using the pathfindR package. Transcription factor binding analysis involved two Enrichr tools: ChEA (2016) and Transcription Factor Perturbations [18]. For both analyses, adjusted P-values (computed using the Benjamini-Hochberg method to correct for multiple comparisons) were exported and plotted in Graphpad Prism. Molecular signatures were analyzed using the MSigDB Hallmark 2020 collection or the Huntington’s Disease Molecular signatures database (HDSigDB) [19].

### Data Visualization

Fold change values from the differential gene expression analysis (DeSeq2) were calculated as described [13], imported into Cytoscape [20] and mapped onto networks (Biogrid Protein-Protein interaction Network [21] or GOCAM Gene Ontology network [22]. To create subnetworks for visualization, we first identified hub proteins that were themselves differentially expressed and also contained multiple differentially expressed genes in their nearest neighbors. Colorization of levels of fold change used a continuous scale from red to blue, with grey representing genes not differentially expressed in the comparison groups. For biological processes, the default color palette was changed such that darker blues are more significant adjusted P values, while lighter greens to white are less significant.

Graphs were made in either R (ggplot2 package) or in Graphpad Prism. Venn diagrams were created using online tools available at the University of Gent, Belgium [23].

### Novel Object Recognition

Our protocol for novel object recognition was adapted from Leger et al. [24]. Eight to ten mice of each genotype were used for this study; these mice were 5-7 months old and were approximately sex balanced (50/50 or 60/40). Mice were acclimated to the testing room and to the experimenter for two sessions (two days). On day 3, mice were individually placed in an empty square chamber and allowed to explore for 5 minutes. Videos were recorded from above the box in ambient light and with the experimenter outside the room. On day 4 (training/association), mice were returned to the chamber to explore two identical objects for ten minutes. On day 5 (24 hours after the day 4 training trial), mice were returned to the chamber which contained one familiar object from the day before, along with one novel object. Ten minutes of object exposure was captured via video. Total distance traveled in the open field and time spent within 2 cm of each object during each trial were analyzed offline in Ethovision. To quantify mouse behavior, we used a discrimination index, which is calculated as (total time at novel object – total time at familiar object) / (total object exploration time). Total object exploration time was calculated as the sum of time spent at both the novel and familiar objects during the trial. Positive values indicate recognition of a novel object.

### Elevated Plus Maze

Eight 6-month-old wild-type (5 female, 3 male) and nine *Tmem184b-*mutant (5 female, 4 male) mice were used for the behavior assay. Mice were placed at the distal end of a closed arm and allowed to walk on an elevated X-shaped platform (50cm from the ground, 35 cm arm length, made of gray non-reflective non-odorous poly-methyl methacrylate (PMMA)) for five minutes. Two of the arms of the platform are enclosed by walls, while two are open. The number of times a mouse enters the open or closed arms is recorded. The ratio of open vs. closed entries, and time spent in open versus closed arms, are calculated as an index of anxiety. Benchpads are used under the open arms to cushion any unexpected falls.

### β-galactosidase Staining and Immunohistochemistry

Adult male Thy1-YFP positive mice [25] were euthanized with CO_2_. The hippocampus was removed and briefly fixed in 4% paraformaldehyde for 5 minutes, followed by submersion in X-gal staining solution (0.1 M phosphate buffer pH 7.5, 5 mM potassium ferricyanide, 5 mM potassium ferrocyanide, 20 mM Tris-HCL, 0.02% NP-40, and 0.01% sodium deoxycholate) for 1.5 to 2 hours. Tissues were rinsed and post-fixed in 4% paraformaldehyde/1X PBS overnight. Fixed tissue was washed with PBS, submerged in 30% sucrose/1X PBS, and embedded in O.C.T. media. Sections of 40 μm thickness were cut on a cryostat and incubated with primary antibodies to Rabbit anti-GFAP (Proteintech 16825-1-AP, used at 1:200) and Rat anti-Cd11b (ThermoFisher 12-0112-82, used at 1:200) overnight at 4℃. Secondaries used were Alexa Fluor 488 anti-rabbit (Invitrogen) and Cy3 anti-Rat (Jackson ImmunoResearch).

### Hippocampus Immunostaining

Mice were perfused with 4% PFA/1X PBS, and whole brains were removed and fixed in 4% PFA overnight. The next day, hippocampi were dissected and rehydrated in 30% sucrose 1XPBS. Brain tissue was embedded in O.C.T. media for cryosectioning (40μm thickness) at - 20°C. Tissue sections were blocked with 1% Tx-100, 4% NGS, 2% BSA, and 1X PBS overnight (4°C). Primary antibodies were Rabbit anti- SHANK1 (NovusBio Cat. #NB300-167, used at 1:150) and Mouse anti-PSD-95 (ThermoFisher Cat. #MA1-046, used at 1:200), incubated overnight (4°C). Samples were washed in 1X PBST (0.1% Tx-100 in 1XPBS). Secondaries were incubated for 1 hour and included Alexa Fluor 488 anti-rabbit (Invitrogen) and Cy3 anti-mouse (Jackson ImmunoResearch). Tissue samples were mounted with DAPI antifade vectashield (VectorLabs Cat. #H-1200).

### Image Acquisition and Analysis

Images were acquired on a Zeiss AXIO Observer.Z1 using a 40x oil objective. Images were taken as z-stacks (12) and analyzed as an orthogonal projection. Raw CZI files were imported into ImageJ (Fiji) and processed (background subtraction, white top hat filtering). Images were then binarized and segmented via watershed. We defined nuclei via DAPI and analyzed the surrounding perinuclear area by expanding the DAPI ROI by 3 μm in all directions. We binarized the PSD-95 and SHANK1 images and counted puncta within the expanded and nuclear ROIs using the Process > Find Maxima function. A macro for automating puncta counting across ROIs is available on GitHub (https://github.com/marthab1/image_analysis). To quantify puncta within the perinuclear region, the puncta within the nuclear ROI (inner) was subtracted from the puncta within the enlarged ROI (outer). We analyzed at least 60 neurons per mouse, averaged these to get counts per cell, and evaluated statistical significance using unpaired t-test with Welch’s correction (Graphpad Prism).

## Results

### TMEM184B promotes expression of synapse assembly genes in the mouse hippocampus

TMEM184B expression is required for maintenance of synaptic structure at neuromuscular junctions in both mouse and fly [11, 9]. We wondered if this role would extend to the hippocampus, a center for learning and memory processing. TMEM184B is highly expressed in neuronal layers of the hippocampus compared to other brain regions, suggesting it may significantly contribute here [26]. We compared gene expression in the hippocampus in wild type and *Tmem184b* gene-trap mutant mice (which have less than 5% mRNA of *TMEM184B* remaining [11, 27]) at 5 months of age (Figure 1 and Additional File 2). Wild type mice showed high clustering in both principal components (Figure 1A), while mutant mice separated well from wild type mice in principal component 1 (PC1). In total, 1153 genes were differentially expressed in *Tmem184b-*mutant hippocampi (Figure 1B-C). When considering those genes with significant hippocampus expression, a group of developmentally important transcripts emerged including SHANK1 (involved in post-synaptic density scaffolding) [28, 29], Neurexin 2 (involved in synaptic assembly and adhesion) [30], and Somatostatin (a marker of subtypes of interneurons) [31].

**Figure 1.**
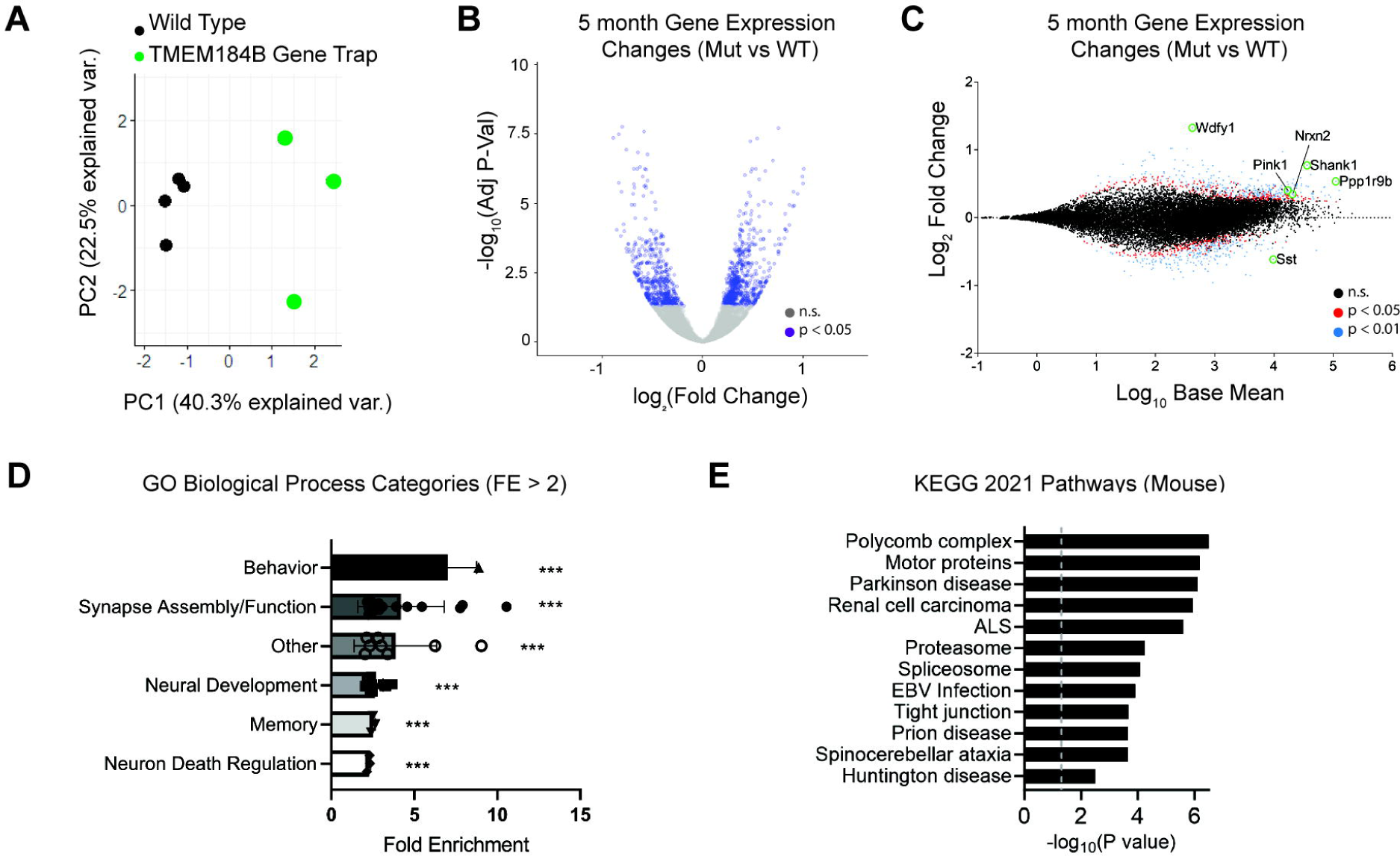
TMEM184B-dependent gene expression alterations in the mouse hippocampus. A, Principal component analysis (PCA) of wild type and mutant samples taken at 5 months of age shows strong clustering of wild types (black circles), and loose clustering of mutant samples (green circles). B, volcano plot showing both up- and down-regulation of genes based on differential expression analysis. Purple indicates adjusted P-values of p < 0.05. C, MA-plot of DESeq2 results. BaseMean is DESeq2’s estimated gene expression across all samples. Red, Adjusted p < 0.05; blue, adjusted p < 0.01. Selected genes of interest are labeled. D, GO Biological Process Fold Enrichment (FE) scores of categories above 2.0 FE. Dots represent individual items in a functional category. P values were calculated as the average p-value for all genes in a category; ***, p < 0.001. E, KEGG Pathways analysis (done in R using the mouse KEGG database and the pathfindR package). Top 12 pathways are shown, ranked by adjusted p-value. Gray dashed line indicates significance threshold (p < 0.05).

To evaluate the biological processes likely affected by TMEM184B in the hippocampus, we used gene ontology analysis to pinpoint processes with significant enrichment of differentially expressed genes in our dataset (Figure 1D and Additional File 3). This showed a large predicted effect on synapse assembly and neural development. Other processes that may be affected in *Tmem184b*-mutant mice include memory/cognition, synapse function, behavior, and neuronal cell death. We performed pathway analysis on these data by querying the mouse KEGG database (Figure 1E) [14–16]. This analysis identified many key cellular pathways such as motor protein and proteasome function that may be disrupted by loss of TMEM184B. Furthermore, it suggested that pathways dysregulated in *Tmem184b-*mutants are enriched for those contributing to neurodegenerative diseases including Parkinson’s disease, ALS, Spinocerebellar ataxia, and Huntington’s Disease. To investigate which cells might be driving these changes, we queried the cell type expression of TMEM184B in the hippocampus. TMEM184B is found primarily in neurons in the hippocampus, with little to no expression in astrocytes or microglia (Supplemental Figure 1 in Additional File 1). Taken together, our analysis of transcriptomic changes in *Tmem184b-*mutant hippocampus predicts significant dysregulation of neuronal function that may be linked to neurological disease.

### Synaptic Protein-Protein Interaction Networks and Biological Processes rely on TMEM184B for proper hippocampal expression

We sought to take a closer look at the relatedness of genes that were differentially expressed in the pathways described above, specifically synapse assembly and neural development. Using Cytoscape [20], we visualized our gene expression changes on the predicted mouse protein-protein interaction network annotated by Biogrid (Figure 2A-B and Supplemental Figure 2). We noticed a significant enrichment of altered transcripts of proteins that are predicted to interact with two key neurodevelopmental proteins: Neuroligin 3 and SHANK1. However, contrary to our expectations, *Tmem184b*-mutant mice show upregulation of transcripts in these networks (shown in circles with pink and red shading). Neuroligin 3 is a cell adhesion molecule that, along with Neurexins, mediate adhesion between pre- and post-synaptic sites. Mutations in Neuroligins are associated with autism spectrum disorders (ASDs) [30, 32]. SHANK1, along with its family members, are critical scaffolds for postsynaptic density proteins and glutamate receptors, and they are similarly implicated in ASDs [28, 29]. The upregulation of these transcripts could be a direct effect of TMEM184B or a compensatory effect following other synaptic disruptions. To evaluate the formation of synapses, we used immunohistochemistry to assess SHANK1 levels alongside those of PSD-95 in the hippocampus. Surprisingly, we found that in areas surrounding neuronal cell bodies, the number of PSD-95 and SHANK1 puncta showed a trend towards a decrease in *Tmem184b*-mutant mice (p = 0.056 for SHANK1) (Supplemental Figure 3 in Additional File 1). These data are consistent with a disruption of synaptic structure in the hippocampus and could suggest that synaptic gene transcripts are upregulated to compensate for lost synapses. Nevertheless, the correct levels of these factors are crucial for maintenance of proper circuit assembly and function; both up- and down-regulation of both proteins is deleterious [33, 34].

**Figure 2.**
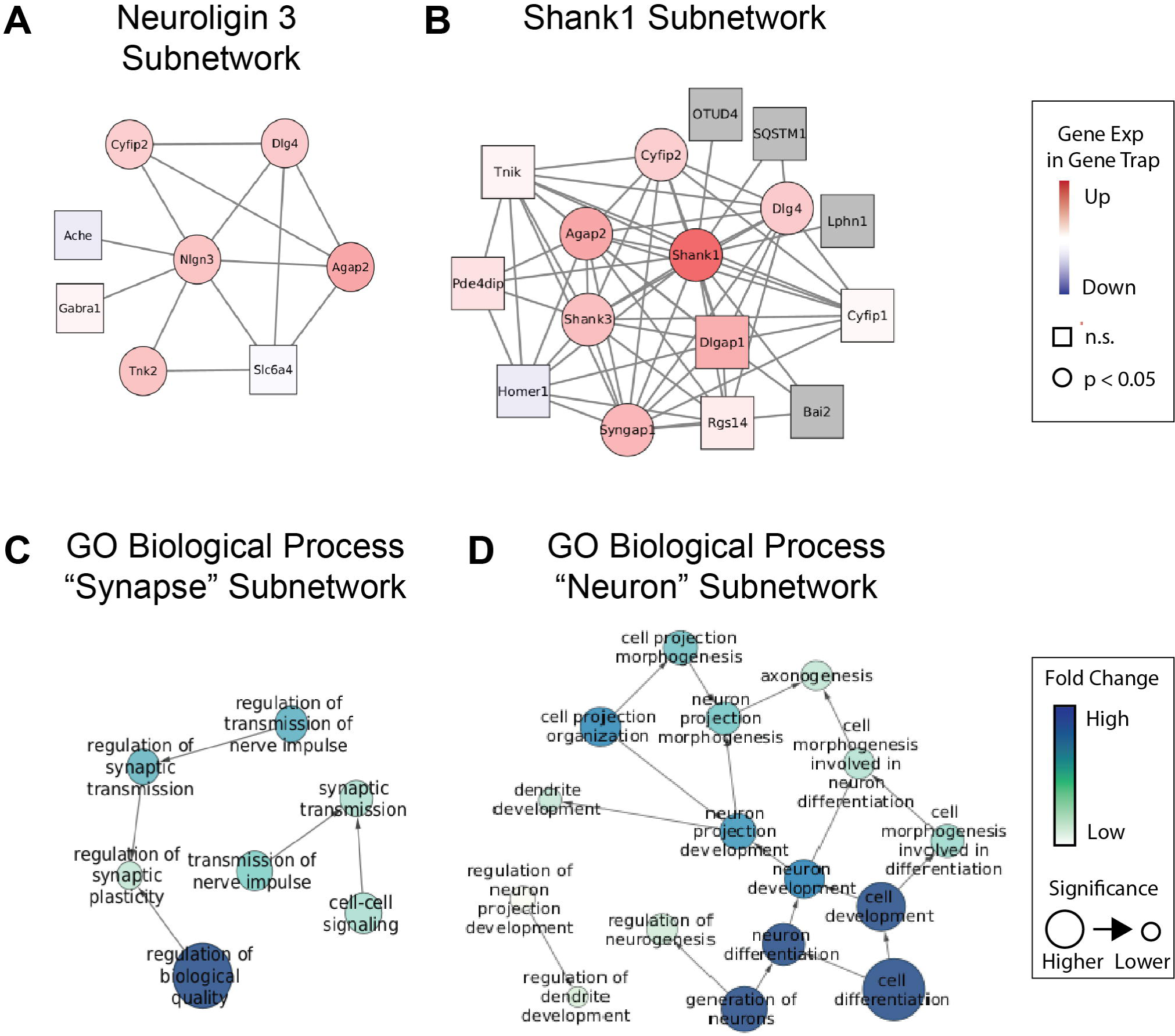
TMEM184B loss disrupts key synaptic gene networks. A-B, mouse protein interaction networks as defined by Biogrid. Node color represents log_2_ of fold change (darker red indicates higher over-expression in *Tmem184b-*mutant; blue shades indicate downregulation in *Tmem184b-*mutant). Circles indicate adjusted P-values where p < 0.05; rectangles indicate p > 0.05. Edge lengths have no significance. C-D, GO Biological Process network maps (from GoCAM) showing terms (gene sets) having significant alteration in *Tmem184b-*mutant mice versus controls. Darker blue indicates stronger fold change, while size of nodes indicates significance (all nodes shown have Adjusted p < 0.05). Edge lengths have no significance. GO Biological Process terms disrupted in *Tmem184b*^+^ hippocampi identified with the search term “synapse” (in C) or “neuron” (in D).

In addition to mapping our data onto known protein-protein interactions, we also mapped our data onto gene ontology (GO) networks and assessed which neuronal or synapse-related processes were predicted to be altered by *TMEM184B* expression (Figure 2C-D). The most differentially expressed GO Biological Processes associated with synapses were in synaptic transmission and plasticity. This prediction is consistent with existing data showing that *Tmem184b* mutation causes hyperexcitability at glutamatergic synapses in *Drosophila* [9]. Within the search term “neuron” these processes were enriched for neurogenesis and projection (axon/dendrite) development. Overall, our analysis suggests that TMEM184B maintains appropriate expression of interconnected networks of synaptic and neurodevelopmental genes in the hippocampus.

### Analysis of Transcription Factors Upstream of Hippocampal Differentially Expressed Genes implicates Wnt signaling alterations

To understand why loss of TMEM184B causes gene expression changes in the hippocampus at 5 months, we performed an *in silico* analysis to identify enriched transcription factor target genes in our DEGs using two platforms. In the first (ChEA 2016), we identified four transcription factors with enriched targets (DMRT1, RARB, ZFP281, and TCF7) (Figure 3A-B). DMRT1 is not expressed in the brain and so is unlikely to have an effect. ZFP281 is a transcriptional repressor that maintains the pluripotency of stem cell populations and is highly expressed [35]. RARB, or retinoic acid receptor beta, is a steroid hormone receptor responsive to retinoic acid (RA) and plays roles in embryonic development. Disruptions to RARB lead to neurodevelopmental delay and are associated with autism spectrum disorder [36]. TCF7 is a canonical Wnt signaling pathway mediator [37]. In a separate analysis, we used the EnrichR platform to examine the overlap between DEGs from our data set with mouse transcription factor manipulations performed by others. We identified many transcription factors whose gene expression alterations significantly resemble that of *Tmem184b*-mutant hippocampus (Figure 3C). Interestingly, multiple associations were identified with transcription factors known to promote Wnt signaling (NEUROD1, TCF3, PLAGL2) [38–40]. This is consistent with prior work suggesting that Wnt pathway dysfunction could participate in TMEM184B-dependent phenotypes [27]. Wnt/β-catenin signaling is critical for hippocampal neurogenesis [41, 42], so this suggests that one effect of *Tmem184b* disruption may be loss of neurons. Other factors, such as NFIA, are known contributors to stem cell maintenance and proliferation. Adult neurogenesis in the hippocampus is required for long-term spatial memory in mice [43]. Taken together, this analysis suggests a possible role for TMEM184B in the generation of hippocampal neurons via promotion of Wnt pathway signaling.

**Figure 3.**
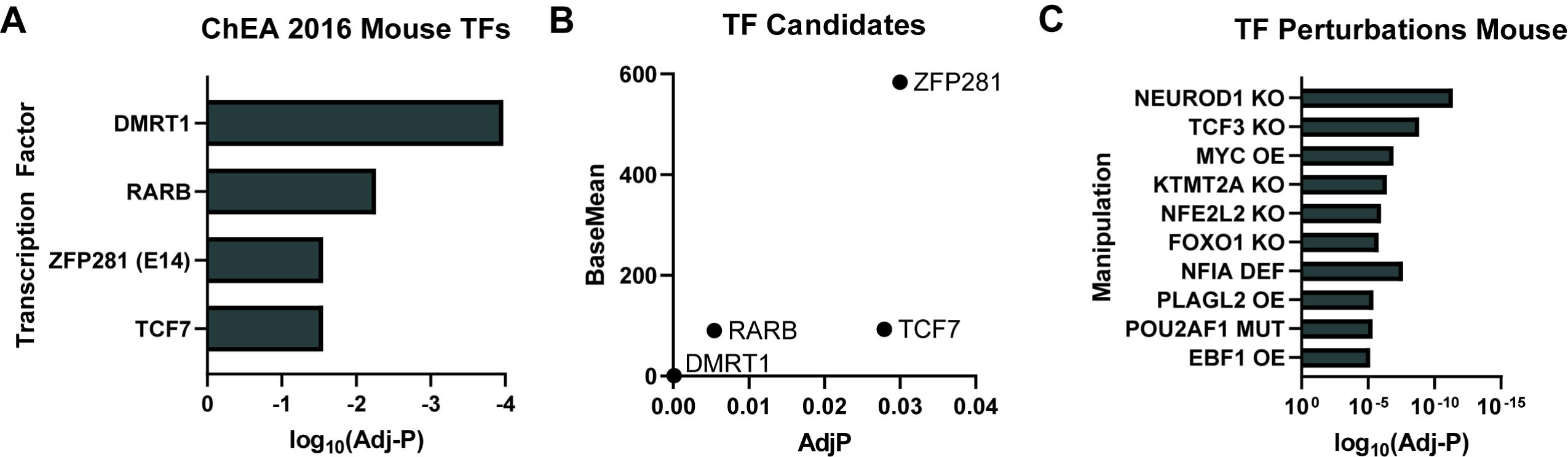
Analysis of Transcription Factors Upstream of Hippocampal DEGs identifies Wnt pathway alterations. A, Chromatin Enrichment Analysis (ChEA) for mouse transcription factors (TFs) with binding in promoters of differentially expressed genes from *Tmem184b*-mutant hippocampi at 5 months of age. Shown are TFs for which enrichment scores were statistically significant (Adjusted P < 0.05). B, Base mean expression in 5 month old hippocampus of each transcription factor in A, graphed against its adjusted P-value. C, top 10 mouse data sets from transcription factor manipulated genetic backgrounds that show significant overlap with TMEM184B differentially expressed genes. Adjusted P values calculated by Enrichr.

### Identification of Common and Unique Aging Signatures in *Tmem184b*-mutant hippocampus

The biggest risk factor for the development of Alzheimer’s Disease is advanced age. To identify unique signatures of TMEM184B disruption that may affect the aging brain, we performed additional RNA sequencing on the hippocampi of older *Tmem184b-*mutant or wild type mice (15 months of age). We then identified genes differentially expressed across aging, genotype, or both. The *TMEM184B* transcript itself was not significantly altered across wild type aging, although it was slightly reduced in older mice (log_2_FC = −0.13; Adjusted P value = 0.074).

After filtering out ribosomal transcripts, a total of 2171 genes were significantly altered in 15-month-old *Tmem184b-*mutant mice compared to age-matched wild types (Additional File 4). Of these, 222 transcripts were altered at both ages (Figure 4A-B and Additional File 5). We identified the most enriched processes in this group using gene ontology (Figure 4C-D). This analysis confirmed the strong effects of TMEM184B loss on synaptic processes, including postsynaptic density organization, glutamate receptor and neuroligin binding activity, and synaptic plasticity. Interestingly, both young and aged *Tmem184b-*mutant mice have a small but significant upregulation of ApoE transcripts (log_2_FC of 0.23 and 0.30, respectively). This comparison confirms that, throughout the aging process, TMEM184B continues to influence synaptic gene expression.

**Figure 4.**
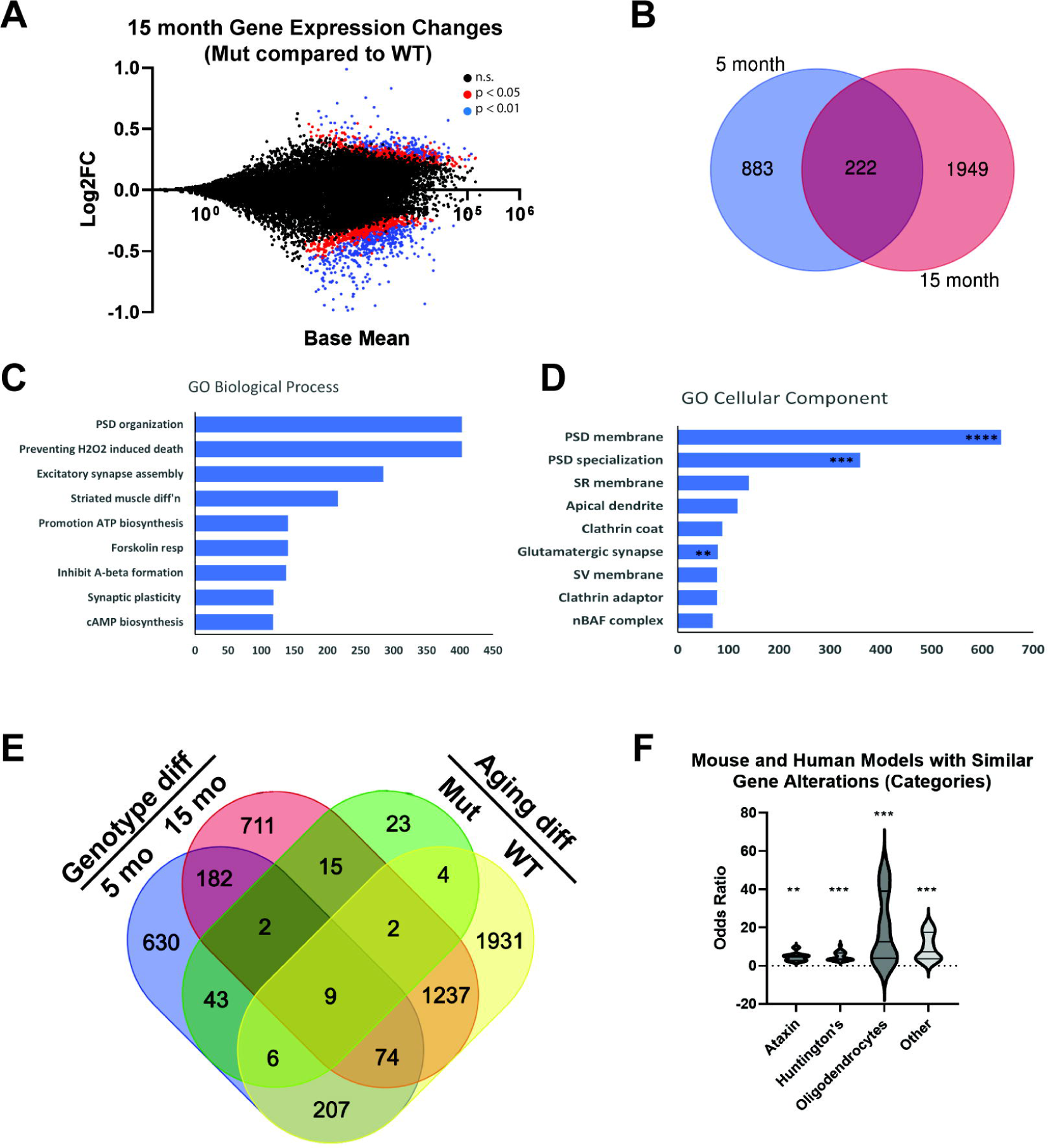
Gene Expression Changes in 15-month *Tmem184b*-mutant vs wild type mice. A, MA plot of gene expression changes between 15-month-old *Tmem184b*-mutant and wild type hippocampus. Red dots show genes with adjusted p < 0.05, while blue dots show adjusted p < 0.01. B, Venn diagram showing overlap between 5 month changes and 15 month genotype-induced changes. C-D, evaluation of enrichment among the 222 overlapping genes using gene ontology for Biological Processes (C) and Cellular Components (D). Shown is Combined Score reported by Enrichr. This score takes into account both P-value and Odds Ratio. Asterisks indicate level of statistical significance (Adjusted P-value, **** <0.0001, *** < 0.001, ** < 0.01). E, Four-way comparison of differentially expressed genes (no ribosomal gene filtering). F, the top 50 gene set categories identified by Molecular Signatures Database (MSigDB) among genes different between 5- and 15-month *Tmem184b-*mutant mice. Abbreviations: HD, Huntington’s Disease model mice; Oligo, oligodendrocytes; Ataxin, Ataxin-1 mutant mice. The full list of molecular signatures is shown in Additional File 7.

If we compare 5-month-old and 15-month-old mutant mice, very few genes are significantly different (104 total) (Figure 4E and Additional File 6). In this group, no biological pathways, molecular functions, or cellular components reached statistical significance (all had adjusted P values > 0.05). This indicates that the changes that have occurred in *Tmem184b-* mutant mice primarily occur by 5 months. Of the 104 genes found to be disrupted, many are disrupted in other mouse models of neurological disease, including two Huntington’s Disease models (R6/2 and Q175) and spinal cerebellar ataxia models caused by disruption of Ataxin-1 (Figure 4F and Additional File 7). In a broader look at the molecular signatures of TMEM184B disruption using the MSigDB Hallmark 2020 collection, we identified two gene sets with statistically significant similarity to ours: TNF-alpha signaling (p = 0.02) and Adipogenesis (p = 0.02).

### Somatostatin, and other neuronal genes, are similarly regulated in both central and peripheral neurons

To evaluate common signatures of TMEM184B-dependent gene expression across disparate types of neurons, we compared the differentially expressed genes (Adjusted p < 0.05) between our 5 month hippocampus data set and one we have previously reported from 6 month adult dorsal root ganglia [10]. Among differentially expressed genes, we identified 12 genes (including *Tmem184b* itself) that were similarly regulated (Figure 5 and Additional File 8). In many cases, the fold change in the hippocampus was somewhat less than that in the DRG, which could reflect a greater percentage of non-neuronal cells in the hippocampus that may dilute the overall fold change. Nevertheless, we noticed that Somatostatin, a neuropeptide used as a marker of GABAergic inhibitory interneurons in the hippocampus (but which marks excitatory, pruriceptive neurons in the DRG), was downregulated in both data sets, suggesting a common means of regulation in these two tissues. One common upregulated gene, Tsukushi (*Tsku*), is a negative regulator of the Wnt pathway [44]. Taken together with our earlier analyses [10], this again implicates a loss of Wnt signaling in TMEM184B-associated phenotypes.

**Figure 5.**
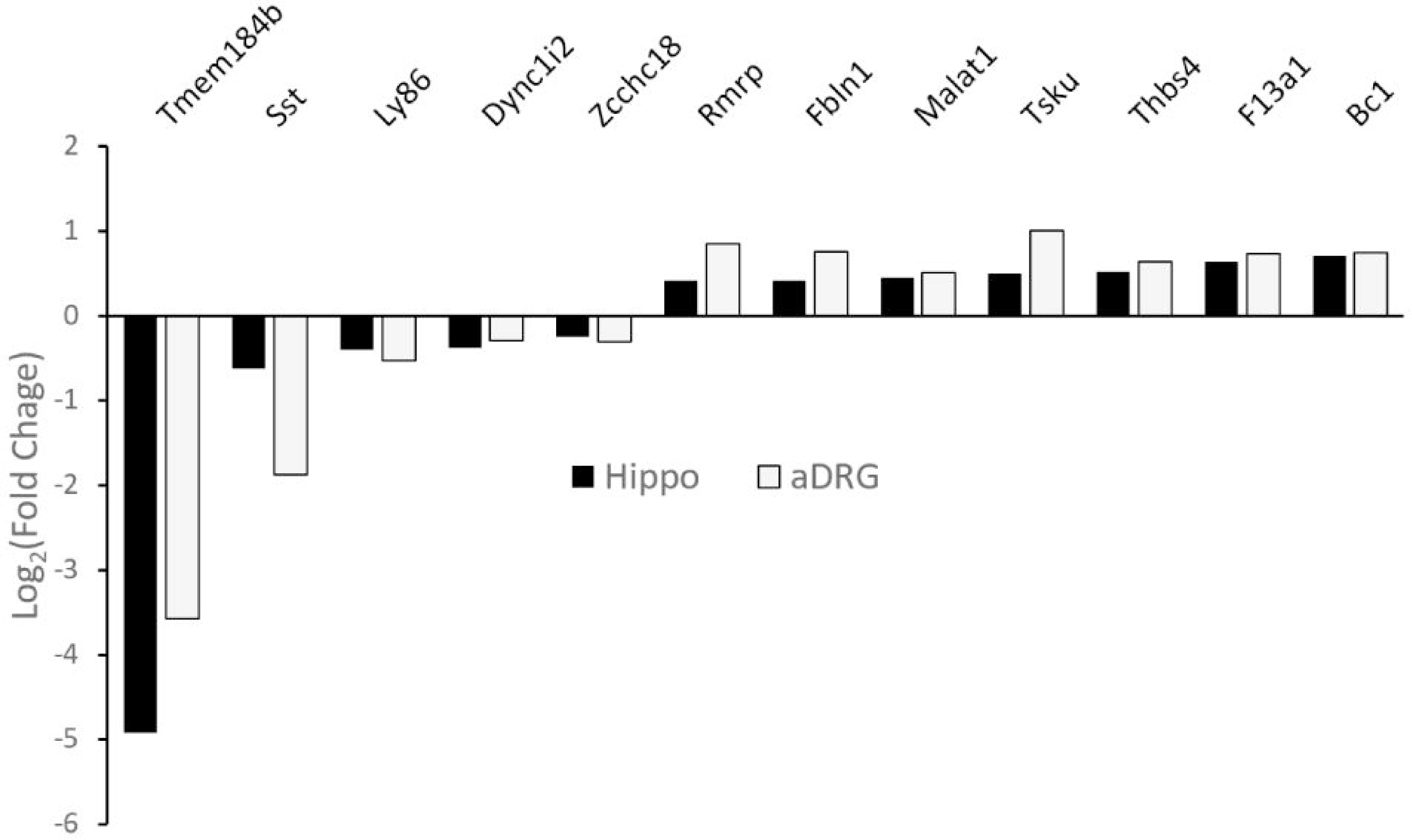
TMEM184B similarly regulates somatostatin and other transcripts in both central and peripheral neurons. Data for DRG is available at NCBI GEO (NCBI GEO GSM4668859) Log_2_FC of genes identified in each data set (hippocampus, black bars; adult DRG, grey bars). A full list of genes differentially regulated in both data sets can be found in Additional File 8.

### TMEM184B loss does not affect object-oriented memory but alters anxiety behaviors in middle aged mice

Finally, we sought to evaluate how these gene expression changes in the hippocampus affect behaviors influenced by this brain region, including anxiety and object-oriented memory. In prior work, we established that six-month-old *Tmem184b-*mutant mice show no deficiencies on the rotarod assay, but show difficulty in an inverted screen test, indicating that some sensorimotor impairment is present [10, 11]. Prior to performing memory assays, we first evaluated mobility of mice in an open field, using the same arena we planned to use for memory testing. We did not observe any significant differences in mobility (total distance traveled) (Figure 6A), indicating that we could use this paradigm and arena for novel object recognition, a classical object-oriented memory paradigm. However, we did not see any difference between wild type and *Tmem184b-*mutant mice in their ability to recognize novel objects (both showed a positive discrimination index, indicating more time at the novel object) (Figure 6B). This indicates that at six months of age, TMEM184B loss does not alter object-oriented memory in this paradigm.

**Figure 6.**
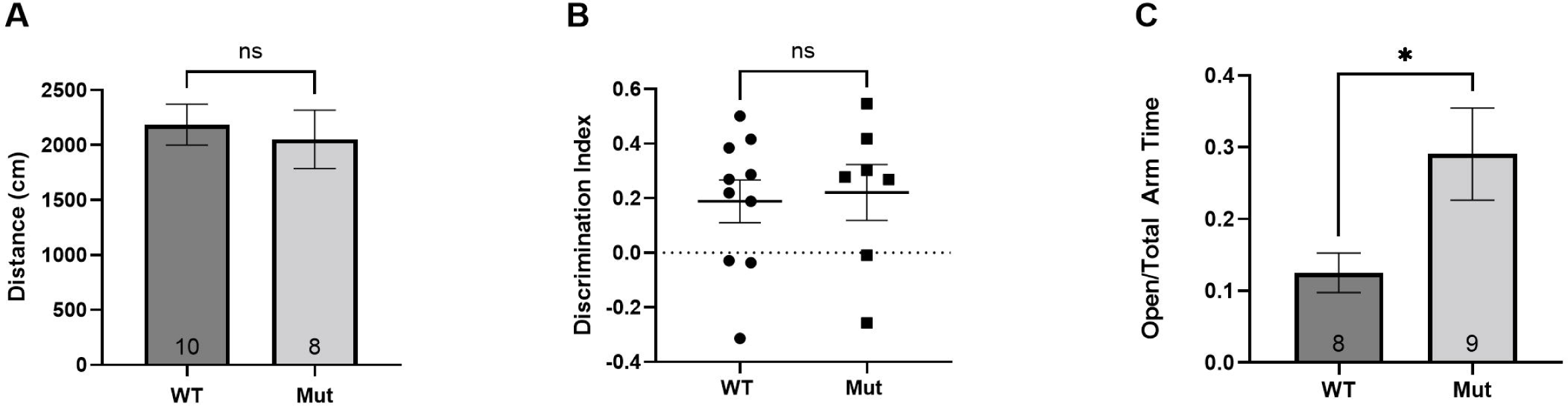
Memory and Anxiety Behaviors in TMEM184B deficient mice. All graphs show mice of both sexes and are approximately sex balanced. **A**, Open field habituation over five minutes for mice 5-7 months of age. Total distance traveled in centimeters. WT, wild type, Mut, *Tmem184b*-gene trap mutant mice. **B**, Memory test at 24 hours using the novel object recognition task. Discrimination index is calculated as described in materials and methods. Each dot shows an individual mouse. N = 10 wild type and 8 mutant. **C**, measurement of anxiety using the elevated plus maze. For all graphs, statistical evaluation used unpaired t-test with Welch’s correction. Numbers in bars (in A and C) indicate sample size. Error bars show standard error of the mean (SEM) for all panels.

Next, we evaluated anxiety using the elevated plus maze. Anxious mice spend more time in closed arms (with walls) than in open arms (without walls). Surprisingly, we identified a decreased anxiety overall in *Tmem184b-*mutant mice when compared to wild types. *Tmem184b*-mutant mice spent considerably more time in the open arms of the maze (Figure 6C). When data is separated by sex, females but not males were found to drive this difference in anxiety behavior (Supplemental Figure 4 in Additional File 1). This indicates that TMEM184B is required for appropriate anxiety levels in females in the elevated plus maze assay.

## Discussion

TMEM184B was predicted in human transcriptome analyses to be associated with deleterious gene expression that could drive the progression of Alzheimer’s dementia [8]. Our experiments sought to evaluate this hypothesis using RNAseq analysis across aging in young and aged *Tmem184b*-mutant and wild type mice. Our results show that TMEM184B has significant effects on synaptic gene expression in the hippocampus in both young and old mice.

Alteration of post-synaptic density gene transcripts occurred in all of our pathway analyses. Proteins in this group, such as SHANK1, SHANK3, and PSD-95/DLG4 contribute to the scaffolding of neurotransmitter receptors [45, 28]. Interestingly, transcripts of all three of these genes are over-expressed in *Tmem184b*-mutant mice. This upregulation of mRNA could reflect a compensation for presynaptic dysfunction, an inappropriate dendritic overgrowth, or a failure to prune exuberant synapses. In immunohistochemical analysis, we observe trends toward reduced numbers of postsynaptic puncta, which may support the compensation model. Further detailed analysis of hippocampal synaptic morphology and physiology will be necessary to parse apart these possibilities.

Somatostatin, a common downregulated gene across both central and peripheral *Tmem184b*-deficient data sets, plays many distinct roles in the nervous system. In dorsal root ganglia, it is a neurotransmitter for a subset of itch-activated sensory neurons [46]. TMEM184B loss in DRG neurons causes disruption of sensation of interleukin-31, a key cytokine involved in atopic dermatitis (eczema) that is detected exclusively by SST*+* neurons [46, 47]. In contrast, in the dentate gyrus, somatostatin reduction occurs in genetic models of temporal lobe epilepsy [48]. Reintroduction of SST in this model reverses the epileptic phenotypes, arguing for a causal role in suppression of seizures. *Drosophila* models of TMEM184B deficiencies show ectopic firing and elevated synaptic calcium, indicating a role in restraining excitability [9]. Thus, phenotypes of SST and TMEM184B overlap substantially; our determination that TMEM184B regulates *Sst* expression in multiple cell types could explain this concordance. It would be of future interest to determine whether restoration of SST in *Tmem184b*-mutant mice could restore normal gene expression and/or behaviors.

Another common theme among our data is the connection between TMEM184B and Wnt pathway regulation. In our prior work, we showed that TMEM184B expression positively influences the expression of many Wnt signaling components, as well as their downstream targets, in developing somatosensory neurons [10]. Here we also find dysregulation of Wnt components in the hippocampus, including the loss of Wnt pathway targets as well as upregulation of Wnt pathway inhibitors such as Tsukushi. Wnt signaling deficiency contributes to AD phenotypes in mouse models [49], and it has also been identified as a key pathway altered in human AD samples [50]. While our data does not directly link TMEM184B to the development of AD pathology, it suggests that TMEM184B dysfunction disrupts synaptic gene regulatory networks and also influences expression of genes with more direct links to AD.

The mechanism of action of the TMEM184B protein is yet unknown. Clues to its role include localization to endosomes and synaptic vesicles, accumulation of multilamellar structures in mutant presynaptic terminals, and additional autophagosomes and lysosomes in skeletal muscle in its absence [11]. Because Wnt signaling occurs within endosomes and contributes to dendritic growth, we hypothesize that a failure of Wnt signaling (perhaps due to a blockade in the endolysosomal pathway) may account for some of the phenotypes we observe.

While we did not complete an exhaustive behavioral analysis, we found that TMEM184B disruption causes reduced anxiety in females in the elevated plus maze paradigm. Reduced anxiety is seen in some models of neuroatypical development including autism spectrum disorders and in mice predisposed to depression [51, 52]. Data from lesion studies, optogenetic manipulations, and immediate early gene expression in the hippocampus support a separation of memory and anxiety control centers in the hippocampus, with the ventral hippocampus primarily responsive to anxiety-inducing stimuli [53, 54].

Genetic associations with anxiety have not been well elucidated. Human dataset analysis has identified a group of 26 risk genes associated with anxiety using transcriptome-wide association (TWAS) [55], but none of these were significantly altered in our dataset. Therefore we suspect that it is an overall network change, rather than any individual gene, that is contributing to synaptic alterations leading to our observation of decreased anxiety in *Tmem184b*-mutant mice, especially in females. We hypothesize that disruptions to neurogenesis (GO term 0050767, 2.14 fold enriched) and synaptic organization (GO term 0050808, 2.9 fold enriched) could contribute to these changes, especially given that hippocampal adult neurogenesis is required for proper anxiety circuit function [56]. Here again, Wnt signaling is known to play a role. During neurogenesis, canonical Wnt signaling (through β-catenin) triggers mouse hippocampal neural stem cells to differentiate towards neuronal fates and integrate into circuits [41, 57]. Thus it is plausible that in the absence of TMEM184B, Wnt signaling pathways show reduced activation, leading to a decrease in adult neurogenesis, a disruption of hippocampal function, and an altered anxiety response; these links must be formally tested in future work. These changes may be accentuated in females compared to males. Indeed, in humans, anxiety-related diagnoses are much more prevalent in females [58]. Our RNAseq experiments were not powered to determine a difference by sex. It will be of interest in the future to explore how TMEM184B-controlled molecular pathways may have sexual dimorphism.

We did not observe object-oriented recognition memory impairment when TMEM184B was disrupted in 6-month-old mice. Future work should evaluate other types of memory in these mice, specifically spatial memory (using the Morris water maze task) or associative learning (using fear conditioning paradigms). It is still formally possible that older mice may show impairments. However, perhaps more likely, we suspect that TMEM184B disruption itself is not causative for memory loss. Instead, we hypothesize that compromising TMEM184B function, in the presence of other susceptibility loci, may increase the likelihood of developing dementia. Given its strong effects on synaptic gene expression networks as well as the disruptions to synaptic morphology and function seen in other studies, it will be of significant future interest to directly evaluate synaptic function via electrical recording in the hippocampus and to evaluate its role in the context of mouse models of AD.

## Author Contributions

E.W., E.L., and H.H. analyzed RNAseq data. E.W. performed mouse behavioral tests. E.W. and C.C-R. performed immunohistochemistry. All authors assembled the figures and wrote the manuscript. M.R.C.B. funded the research.

## Availability of Data and Materials

The data sets supporting the conclusions of this article are available in the NCBI GEO repository [GSE204831; https://www.ncbi.nlm.nih.gov/geo/query/acc.cgi?acc=GSE204831]. All custom-written code in R for RNAseq analysis is available at GitHub [https://github.com/eriklarsen4/ggplot-scripts.git and https://github.com/marthab1/image_analysis]. Dorsal root ganglion data to which we compared our hippocampal RNAseq data can be found on NCBI GEO (GSE154316).

## Acknowledgements

The authors would like to thank Ms. Rachel Dinh for help with initial analysis, Ms. Tiffany Cho for genotyping assistance and behavior consultation, Mr. Matt Schmit for behavior consultation, perfusion help and Ethovision training, Dr. Tally M. Largent-Milnes for behavior consultation, and Dr. Rajesh Khanna for use of the elevated plus maze. We would like to thank Dr. Rui Chang for consultations early in the development of this project.

## Funding

This work was funded by the NIH (NS105680) Alzheimer’s Disease and Related Dementia supplement to M.R.C.B.

## Conflict of Interest

The authors do not report any conflicts of interest pertaining to the content of this manuscript.

## Ethics Approval and Consent to Participate

All animal experiments were approved by the University of Arizona Institutional Animal Care and Use Committee (IACUC) Protocol 17-216. All methods were carried out in accordance with relevant guidelines and regulations regarding the ethical treatment of animals. All methods are reported in accordance with ARRIVE guidelines for the reporting of animal experiments.

## Consent for Publication

Not applicable.

## Notes

### Competing Interest Statement

The authors have declared no competing interest.

### Summary of Updates

Author list, methods, and supplementary files updated.

