## Supplemental Files 1-8 for "Transmembrane protein 184B (TMEM184B) promotes expression of synaptic gene networks in the mouse hippocampus": Additional File 1 - SuppFigs 1-4.docx

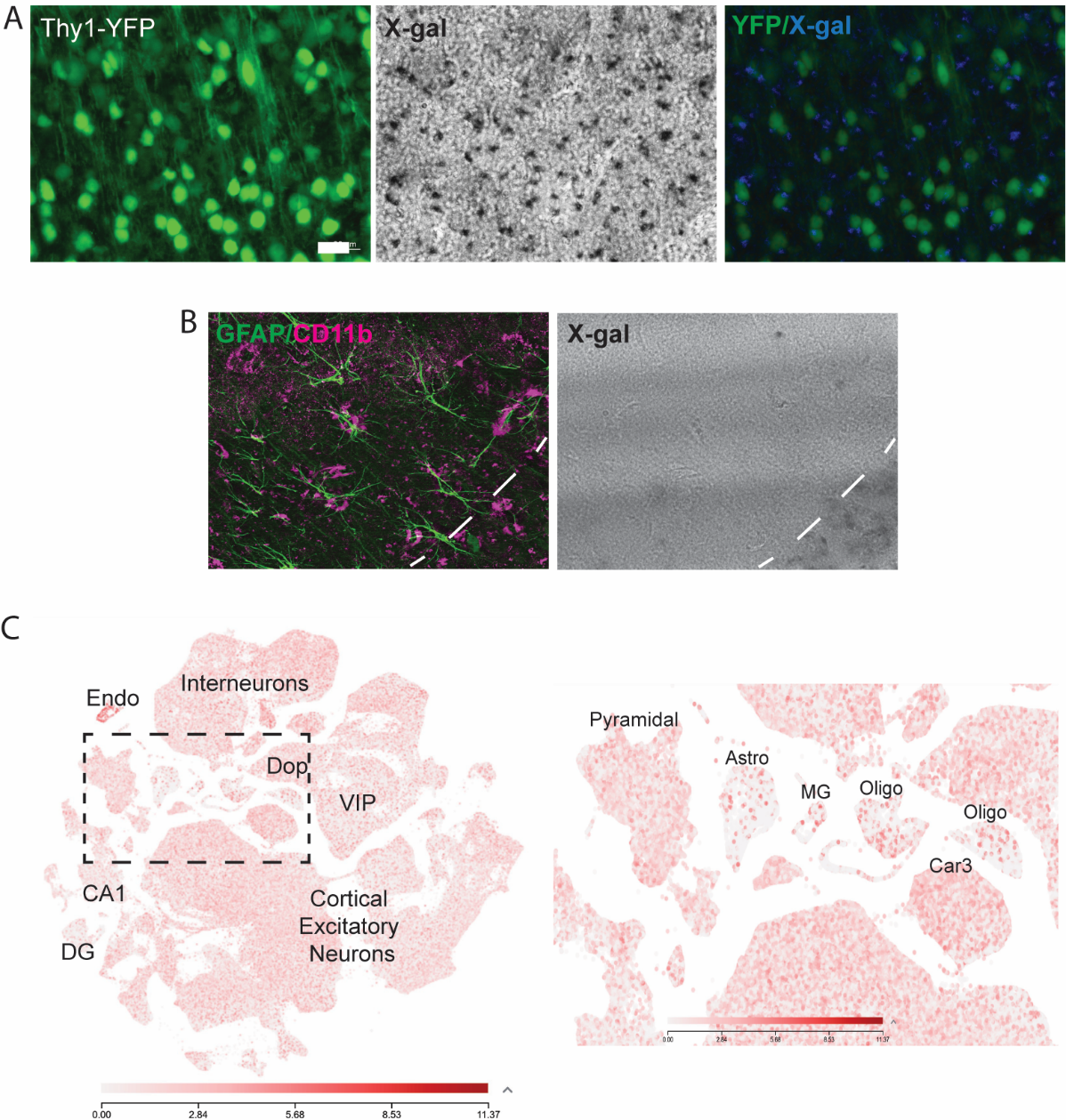

**Supplemental Figure 1. *TMEM184B* is enriched in neurons in the hippocampus.** In A-B, X-gal staining indicates b-galactosidase expression which is driven by the Tmem184b endogenous promoter in heterozygous gene-trap (mutant) mice. A, hippocampal molecular layer labeled with Thy1-GFP (neurons) and X-gal (dark precipitate). Merged image (right) shows similar (overlapping or juxtaposed) labeling of hippocampal neurons. B, hippocampal sections immunostained to highlight astrocytes (GFAP, green), microglia (CD11b, magenta), or X-gal. The bottom right corner (separated by a dotted line) shows the neuronal cell bodies (from A) . Very little x-gal precipitate is found in astrocytes or microglia. C, Allen Brain Transcriptome Atlas showing *TMEM184B* expression from single cell RNAseq dataset of the adult mouse hippocampus and cortex. Right panel is an enlargement of the left panel highlighting expression levels in astrocytes (Astro), microglia (MG), and oligodendrocytes (Oligo). *TMEM184B* transcripts are found in low but consistent levels throughout neuronal populations in the hippocampus and cortex, but they are comparatively very low or absent in glial cell types. Other abbreviations: Endo (endothelial cells), DG (dentate gyrus). Other cell types are as labeled.

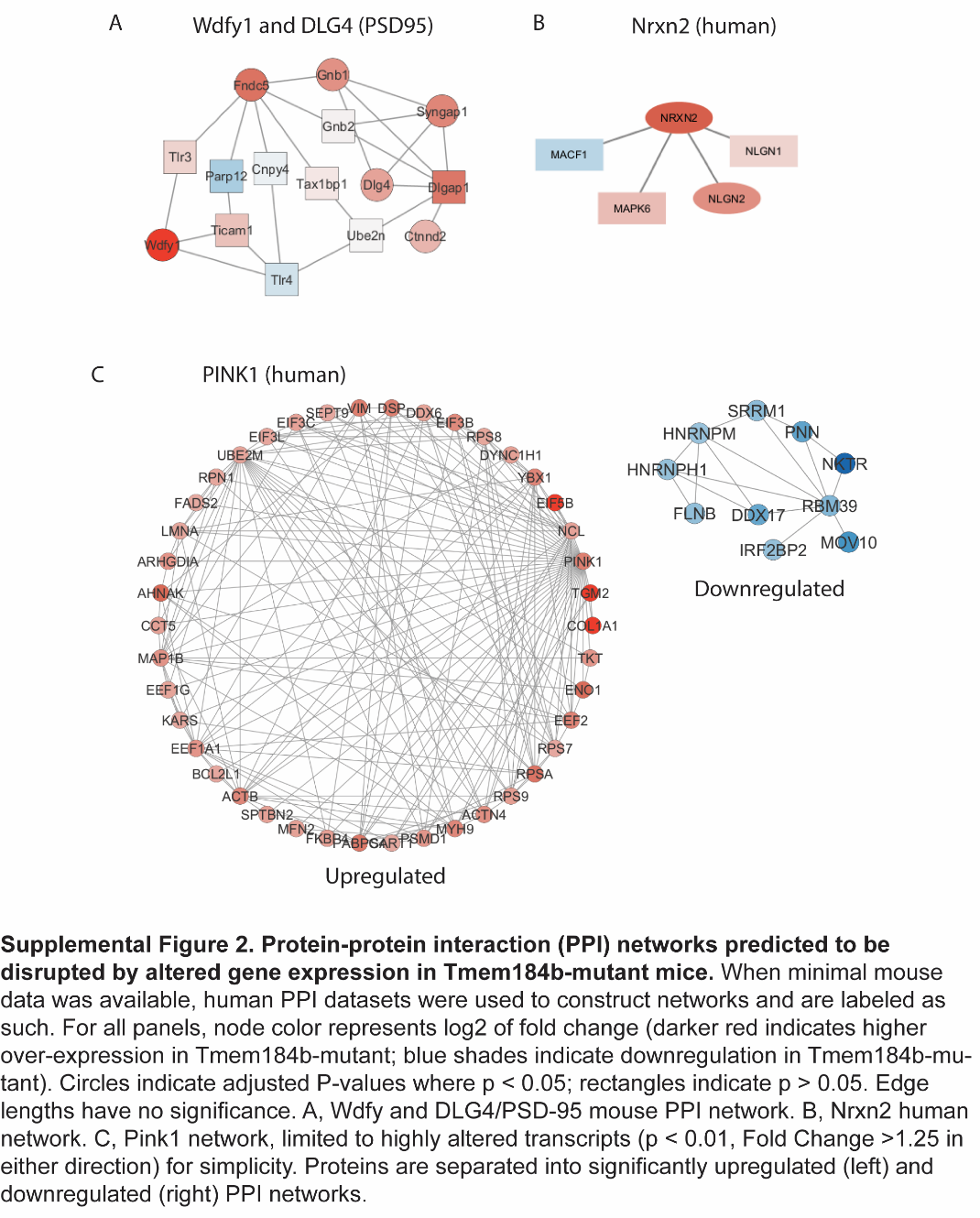

**Supplemental Figure 2. Protein-protein interaction (PPI) networks predicted to be disrupted by altered gene expression in *Tmem184b*-mutant mice.** When minimal mouse data was available, human PPI datasets were used to construct networks and are labeled as such. For all panels, node color represents log_2_ of fold change (darker red indicates higher over-expression in *Tmem184b-*mutant; blue shades indicate downregulation in *Tmem184b-*mutant). Circles indicate adjusted P-values where p < 0.05; rectangles indicate p > 0.05. Edge lengths have no significance. A, Wdfy and DLG4/PSD-95 mouse PPI network. B, Nrxn2 human network. C, Pink1 network, limited to significantly altered transcripts (p < 0.05, Fold Change >1.25 in either direction) for simplicity. Proteins are separated into significantly upregulated (left) and downregulated (right) PPI networks.

**SHANK1**

**WT**

**Mutant**

**WT**

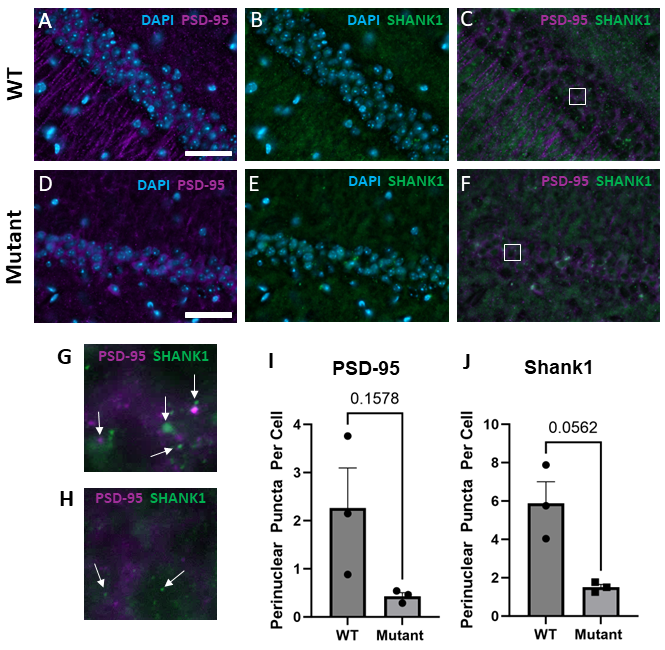

**Supplemental Figure 3.** Assessment of PSD-95 and SHANK1 protein expression in the hippocampus of *Tmem184b*-mutant mice. A-F, Representative images of PSD-95 (magenta), SHANK1 (green), and DAPI (blue) expression in hippocampal sections. Scale bars = 40μm. A-C, wild-type; D-F, *Tmem184b-*mutant. G-H, Enlarged, cropped images of wild-type and *Tmem184b-*mutant puncta from C and F, respectively. White boxes in C and F outline the specific regions enlarged. Quantification of perinuclear PSD-95 (I) and SHANK1 (J) expression in hippocampal sections. Statistical evaluations between groups used unpaired t-test with Welch’s correction for unequal variance. Error bars show standard error of the mean (SEM). P values are listed above each comparison.

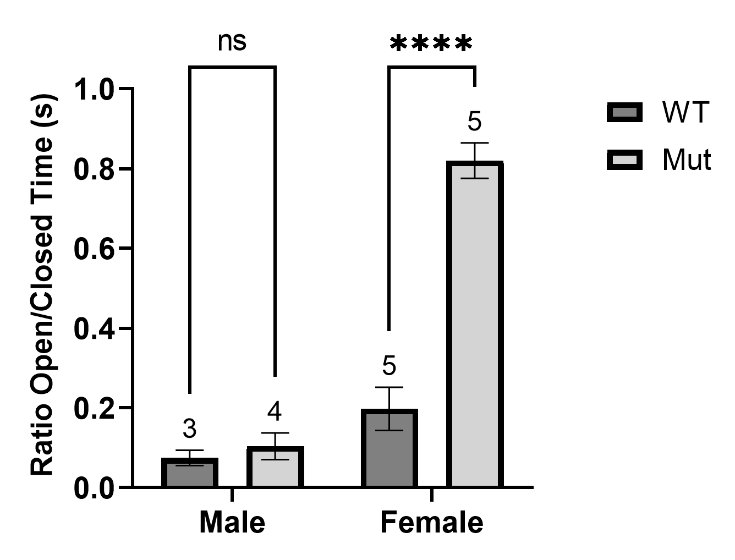

**Supplemental Figure 4. Female *Tmem184b*-mutant mice show reduced anxiety in the elevated plus maze**. Statistical evaluations between groups used unpaired t-test with Welch’s correction. Numbers above bars indicate sample size. Error bars show standard error of the mean (SEM). Adjusted P-value, ****, p < 0.0001.
